# Emergent remote stress-focusing drive submucosal collagen fiber remodeling in intestinal anastomotic surgery

**DOI:** 10.1101/2022.10.31.514540

**Authors:** Brian P. Fleischer, Nhung Nguyen, Anshuman Pal, Julius Kim, Kameel Khabaz, Alkiviadis Tsamis, Efi Efrati, Thomas Witten, John C. Alverdy, Luka Pocivavsek

## Abstract

Following resection of a diseased segment of intestine, a reconnection (anastomotic) geometry is chosen to reduce postoperative stress and optimize outcomes. As proper healing of an intestinal anastomosis is strongly affected by its mechanobiology, much attention has been devoted to the conical structures formed along the suture lines, where stress-focusing is expected. However, geometric considerations reveal that in addition to the obvious loci of stress-focusing, additional remote locations of stress-focusing may form. We identify conical structures that inevitably form within regions of otherwise uninterrupted tissue. In this work we use geometric analysis, finite element modeling (FEM), and in-vivo experiments to investigate these emergent stress-focusing structures, their mechanical stresses, and the resulting submucosal collagen fiber re-orientation, as these naturally arise in the side-to-side small bowel anastomosis (SBA), the most common configuration performed in patients. FEM predicts the appearance of remote high-stress regions. Allowing for tissue remodeling, our simulations also predict an increased dispersion of submucosal collagen fibers in these regions. In-vivo experiments performed on ten-week-old male C57BL/6 mice assigned the creation of side-to-side SBA or sham-laparotomy corroborate this result. Anastomoses were analyzed at sacrifice on post-operative day (POD) 14 and 88 with histologic-sectioning, staining, high magnification imaging, and submucosal collagen fiber orientation (*κ*) mapping. The mean and variance of *κ*, a measure of collagen fiber dispersion, at POD-14 far from the anastomosis show similar values to those obtained for sham-operated mice, while the FEM-predicted loci of stress-focusing display statistically significant higher values. The values at POD-88 at all loci show no statistically-significant difference, and agree with those of the sham-operated mice at POD-14.

## Introduction

Reduced to topologic structures all vertebrates are tori: the gastrointestinal track and its many invaginations including the abdominal organs, lungs, and glands form one torus while the cardiovascular system through its extensive branching patterns and anastomosis forms a second high order torus (**? ? ?**) (6). Given the fundamental topology of a metazoan body plan, the anatomy of many organ systems geometrically appear as cylindrical structures. Surgery on the gastrointestinal track and the cardiovascular system is therefore mathematically reducible to changing the underlying topology and geometry of these tissues. Topological changes occur in settings of bypass operations be it intestinal or vascular where the surgeon creates additional loops in the network thereby changing the topology. Geometric changes occur anytime a portion of the network is interrupted and re-connected. Changes in geometry are therefor localized to regions of surgical anastomosis or re-connection and can occur without changes in topology. The central hypotheses of this paper is 1. surgical anastomoses change the local tissue geometry while preserving its topology, 2. these changes in tissue geometry are sources of mechanical stress, and 3. the tissue locally responds to this non-homeostatic stress by changing its structure thereby reducing the stress. By linking geometry to surgical anatomy and surgical anatomy to tissue mechanics, surgeons will be able to purposefully impact anastomotic shape allowing direct translation of knowledge between geometry, mechanical stress, and biology to the patient for optimized anastomotic patency.

### Anastomotic Geometry

The ultimate success of a surgical anastomosis or reconstruction relies on many factors. Since Alexis Carrel pioneered open vascular surgery using direct suturing of hemostatically controlled unpressurized arteries and veins (**?**) and William Halsted revolutionized gastrointestinal surgery through his discovery that the collagen-rich submucosa is the load-bearing layer of the intestinal wall (2), surgeons have sought technical solutions to optimize the outcomes of vascular and gastrointestinal reconstructions (**?**). Despite a century of practice, neither the GI nor vascular anastomosis is perfected to the point that it reliably heals without the two dreaded complications of leakage (anastomotic dehiscence) or stricture (narrowing). Surgeons have largely manipulated the two components of the anastomosis under their direct control: geometry and mechanics (stress).

Focusing on geometry, the fundamental anatomic building block of any anastomosis is a circular cut generating an open cylindrical edge (figure 1a and b) or linear and circular cuts anchored by two vertices (figure 1c and d) generating a geometric perturbation of the native cylindrical geometry (**? ?**). The variety of anastomotic geometries created throughout the last century (figure 1) relies on understanding the native changes in the baseline cylindrical geometry of tissue along its length. First and foremost, both in the GI track and vascular system, different portions of the tubular network have different calibers or diameters. In the GI track, the tissue can be divided into four separate cylindrical structures of different diameters which in decreasing size are the stomach, the large intestine (colon), the small intestine, and the esophagus. As such, transection of different sized cylinders inevitably leads to the fundamental surgical problem of anastomosing two edges (circular or linear) of mismatched length. Figure 1 outlines four canonical solutions to anastomosing two cut cylinders (1–3). In the case of equally matched diameters, such as occurs in small bowel resections, a simple end-to-end length-matched anastomosis can be performed (figure 1a). It is well recognized in the surgical community that such anastomoses have very low complications rates. Indeed, the only source of geometrical incompatibility in this case would be locally along the suture line given the discreet method of mechanical fastening used: suturing or stapling. We will not explore the impact of this local effect further in this paper. In the case of mismatched edge lengths three solutions exist: reducing the mismatch by partially closing the larger cylinder (Billroth 1 anastomosis, figure 1b), closing one cylinder and making a linear cut that matches the circumference of the second transected cylinder (end-to-side anastomosis, figure 1c), or simply closing both cylinders and making linear cuts of equal length that are then fastened together (side-to-side anastomosis, figure 1d). Surgeons appreciate that certain instabilities are inherent in the non-native (figure 1b-d) anastomoses; this is supported by the plethora of commonly placed stabilizing sutures in these anastomoses: the often discussed ‘anti-obstruction stitch’ in side-to-side anastomosis or the purse-string suture placed at the angle of sorrow or ‘Coffin Corner’. Largely driven by the advent of linear stapling devices and the revolutions in laparoscopic and robotic abdominal surgery, the side-to-side (figure 1d) is the most widely performed GI anastomosis, as such the majority of our paper will focus on its geometry (figure 2).

**Fig. 1.**
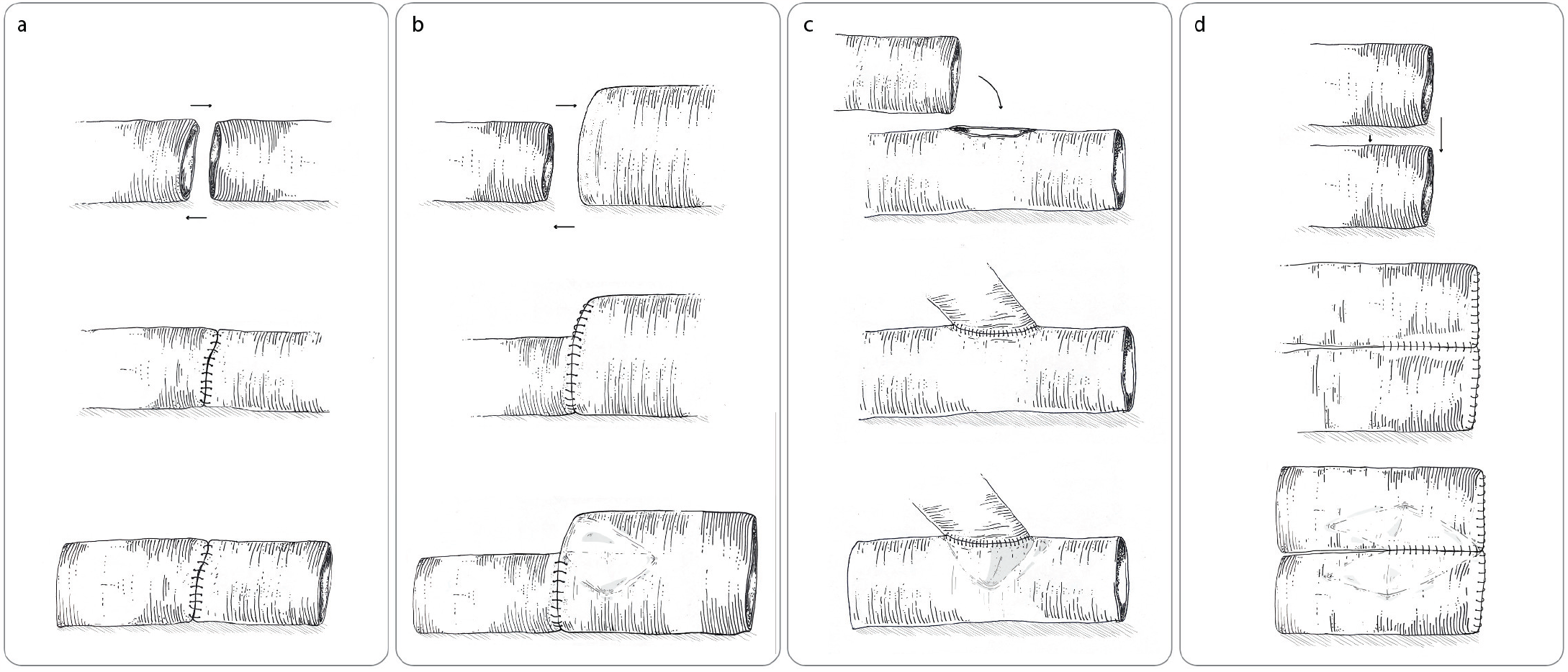
Illustrations of common anastomotic geometries (columns) highlighting their geometric features (top row shows pre-sutured geometry of two limbs, middle row shows un-pressurized sutured geometry, and bottom row shows pressurized sutured geometry with shading indicating regions of conical singularities). (a) End-to-end: this geometry possibly best mimics the preoperative geometry seen in native anatomy, yet it is limited to similarly sized limbs and typically requires manual suturing. No conical singularities appear in a perfectly matched end-to-end. (b) Billroth I: size (circumference) mismatched end-to-end. Here the diameter discrepancy between the two limbs is absorbed by a sutured fold which ends in the well-described angle of Sorrow. In the collapsed state, the Billroth I anastomosis does not seem to have any singularities. Under weak pressurization, strong conical singularities appear in the larger limb. (c) End-to-side: generates a complex region of conical singularities. (d) Side-to-side: designed to avoid the known problems of Billroth I and end-to-side, is also rich in conical singularities. Figure 9 in Appendix B shows FEM simulations of the above anastomotic geometries and highlights the locations of stress-focused conical singularities.

**Fig. 2.**
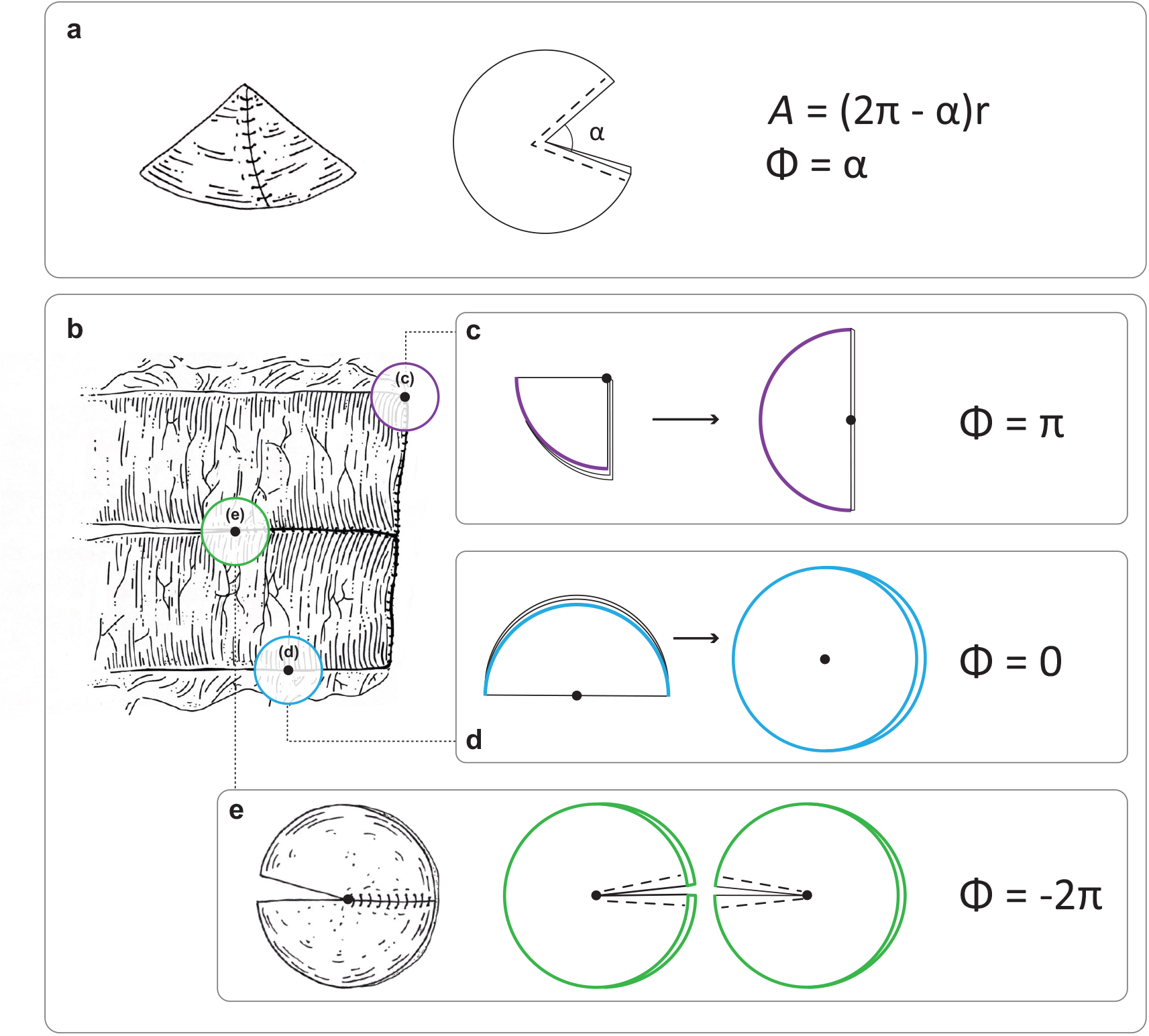
The geometry of conical points in flat (Euclidean) surfaces is defined by the normalized difference between the surface area of all the points whose distance from the singular point is *r* and the area of a flat circle of similar radius, 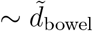 . This formula can be intuitively interpreted as the angular opening of the wedge of material removed to form the singular structure (a). In a side-to-side anastomosis (b), three types of conical singularities appear. (c) Two regular cones with a positive charge (Φ *>* 0) formed by removing a wedge of material occur at the two folded over portions of the suture line closing off the common channel. (d) Internal conical singularities with vanishing charge (Φ = 0) termed *d*-cones occurring in the bowel wall are the main focus of this paper. (e) Singular conical structure with negative charge (Φ *<* 0) called *e*-cones with a saddle like structure occurs at the vertex of the linear staple line connecting the two pieces of bowel.

Prior work from our group explored the geometry of a simple transverse closure or the Heineke-Mikulicz (HM) strictureplasty (9). This particular operation is not commonly performed outside of surgery for certain pathologies such as pyloric stenosis or Crohn’s disease. However, it allowed us to lay down the geometric framework of conical singularities in intestinal reconstructions. Singular conical structures in otherwise flat surfaces can be classified using a geometric charge called the integrated Gaussian curvature (which in the case of singular structures resides at a single point). There are several distinct ways to define the Gaussian curvature (**?**), yet in what follows we only need to consider the special case of point-like Gaussian curvature in otherwise flat (Euclidean) surfaces. In this case, we can identify the integrated Gaussian curvature with the normalized difference between the surface area of all the points whose distance from the singular point is *r* and the area of a flat circle of similar radius, *A*_0_ = *πr*^2^:

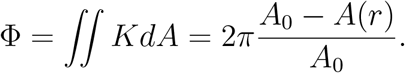

This formula can be intuitively interpreted as the angular opening of the wedge of material removed or added to form the singular structure. In cases where material is added rather than removed the wedge angle is negative, as is the integrated Gaussian curvature. See Figure 2a and Appendix A for more details.

Singular conical structures associated with a positive charge (Φ *>* 0) are formed by removing a wedge of material and are called regular cones. Singular conical structures with negative charge (Φ *<* 0) are called *e*-cones (where ‘*e*’ stands for excess material), and form saddle like structures. Conical singularities with vanishing charge (Φ = 0) are termed *d*-cones (where ‘*d*’ stands for developable, i.e. made from a flat sheet) (**?**). Figure 2b shows three points in side-to-side anastomosis associated with positive, negative, and vanishing values of Φ. Surgically two of the points are well-known to be sources of possible leak: the regular cone (point c. in Figure 2b.) is surgically termed the anastomotic dog-ear and often this region of the anastomosis is reinforced with extra sutures. The *e*-cone (point e in Figure 2b.) is clinically the anastomotic vertex or ‘crotch’, and again extra sutures are often placed in its vicinity to prevent leak or kinking in this area. The location of the *d*-cone has received very little attention clinically.

The tissue reduction or addition that is associated with the creation of regular cones and *e*-cones along the suture line results in macroscopic tissue re-orientation. The remote *d*-cones appear in order to reconcile this re-orientation with the underlying cylindrical geometry. In many instances the unpressurized anastomosed tissue will show no evidence of these remote stress-focusing structures, because the tissue can gently bend to accommodate the weak geometrically imposed incompatibilities in the metric. Once pressurized to allow for luminal content flow, the limbs adopt their cylindrical form away from the suture line and the remote *d*-cones appear. In this paper, we study in detail the geometry of the *d*-cone in the side-to-side anastomosis.

### Anastomotic Mechanics and Remodeling

Carrel in his seminal work on vascular surgery published in 1912 used words such as ‘gentle’, ‘force’, and ‘stress’ dozens of times(**?**); this laid the bedrock of technical principles underpinning modern cardiovascular and transplant surgery where tissue handling is of paramount importance. Likewise, Halsted in 1887 in his seminal paper on the small bowel anastomosis repeatedly writes phrases such as “sufficiently strong to hold a stitch” and in referring to the sub-mucosa he writes “a delicate thread of this tissue is very much stronger and better able to hold a stitch than is a coarse shred of the entire thickness of the muscular and serous coats” and refers to the “resistance furnished by the submucosa” (2). These early and pioneering publications on surgical technique in vascular and GI surgery highlight the importance mechanical stress plays in anastomotic stability (**? ?**). Geometry (i.e., anatomy) influences mechanical stress in rich and fundamental ways (**?**), especially in objects with curvature such as cylindrical arteries, veins, and hollow viscus. Geometrically, thin cylindrical shells can bend more easily than stretch.(**?**) Singular conical defects, *e*-cones at the vertices flanked by *d*-cones, arise to allow the optimal compromise between these bending and stretching modes of deformation.

Surgeons can in large part manipulate the geometry of their anastomosis, guided by two fundamental phenomenologically-derived surgical principles: maintain tissue integrity and strive for a ‘tension-free’ suture line (4, 5). The former confines any viable anastomotic geometry to a class of tissue preserving shapes; meaning any combination of cutting and stitching is allowed however removal of tissue such as beveling beyond disease margins is often not. The latter imposes an even stronger condition on the suture line: length preservation or isometry; suture lines are not to be stretched. The field of surgery suffers from a fundamental lack of understanding of the geometry and mechanics underlying these experientially derived principles. Moreover, the postoperative geometry is often hidden behind non-stress bearing tissue layers, and assumes its true normal functioning form only under internal pressure and content flow. Advanced surgical devices such as staplers and robots have been engineered using empirical principles without a proper scientific basis. Commonly performed operations and hence complications (leaks, strictures, etc) still occur even when performed by expertly trained high volume surgeons operating in the most prestigious institutions. Therefore an in-depth analysis of intestinal anastomotic construction is needed beyond those that have traditionally been ascribed to be causative to complications (ischemia, tension, technique) despite the lack of evidence.

Thin shells bend more easily than they stretch (7, 8), and easily buckle under in plane compressive loads (8). Mechanically modeling the intestine or blood vessel as a thin cylindrical shell that easily buckles was successfully used to predict the typical gut looping morphogenesis observed across many species (6) and well-known arterial tortuosity encountered with aging (**?**). Pure bending deformations with zero in-plane stretching are termed isometries. However, such isometries are not always possible; in lieu of an available isometry the equilibrium configuration will balance the bending and stretching energy contributions. In some cases the optimal compromise between the stretching and bending deformations results in geometrically singular conical structures that focus stress (7, 8).

Previous work from our group has elucidated the critical role that submucosal collagen fibers play in anastomotic healing (12–14); breakdown of these fibers leads to anastomotic leak. In fiberreinforced composites, fiber orientation plays a strong role in setting the global and local mechanics of the system (15, 16). Bowel is living matter endowed with the capacity to locally change tissue structure in response to stimuli. We lack a general understanding on how surgical anastomotic shape impacts local tissue biomechanics and response to injury. While many strong perturbations such as local injury, ischemia, and exposure to collagenolytic bacteria overwhelm the tissue’s ability to respond and cause submucosal collagen fiber breakdown (12, 14), we hypothesize that “softer-modes” of collagen fiber remodeling arise in peri-anastomotic regions with non-native stresses.

**Significance Statement**

Side-to-side anastomosis is the most commonly used geometry for bowel anastomosis, and is generally believed to be the least stressed anastomosis of diamater-mismatched tubular limbs. Geometric considerations and finite element modeling reveal that when the anastomosed tissue is pressurized from its closed rest state to allow for content flow, stress-focusing conical-structures arise in otherwise uninterrupted tissue, away from the suture line. In-vivo study using mouse model reveal significant collagen-fiber remodeling in these stress focusing locations. These results guide….

### Concluding Introduction

In this work we investigate the interaction between intestinal anastomotic geometry, mechanical stress, and collagen fiber distribution and remodeling in side-to-side small bowel anastomosis. We first establish, through FEM, the appearance of *d*-cone type singular structures away from the suture line under pressurization of side-to-side anastomosis. We prove that the obtained *d*-cones are qualitatively and quantitatively crumpling-like structures akin to *d*-cones obtained and studied in less constrained environments (7, 8, 17, 18). Numerically coupling the focussed stresses to tissue remodeling results in increased dispersion of submucosal collagen fibers in the vicinity of the formed *d*-cones. We subsequently design and perform the first ever clinically comparable side-to-side small bowel anastomosis mouse model and show that submucosal collagen fiber away from the suture-line remodels in regions that correspond to the location of *d*-cones in our geometric and finite element analysis.

## Results and Discussion

### Anastomotic Geometry

We simulate the bowel wall using a single layer Ogden-Gasser-Holzapfel (OGH) model, with the material properties adopted from previous work on intestinal mechanics (15). For simplicity and efficiency, we model only one of the limbs of the anastomosis and focus on the formed *e*-cone region and the cylindrical portion leading to it. Further details of material properties and boundary conditions are provided in the Methods section.

When a side-to-side anastomosis is newly created it does not spontaneously open, but remains in a closed configuration, with the longitudinal section in each of the limbs remaining straight and shut. This is especially true when the anastomosis is stapled and adjacent walls are forced to remain tangent to one another. For a functional anastomosis, the intra-luminal pressure within the GI tract provides the needed force to open the anastomosis. When a similar luminal pressure is applied to the simulated anastomosis it opens, and FEM simulations show the existence of locations that exhibit geometric-focusing and high stress. Figure 3 shows a representative deformed geometry of one limb under pressure and the common channel opened to its maximum extent with diameter *d*_*max*_.

**Fig. 3.**
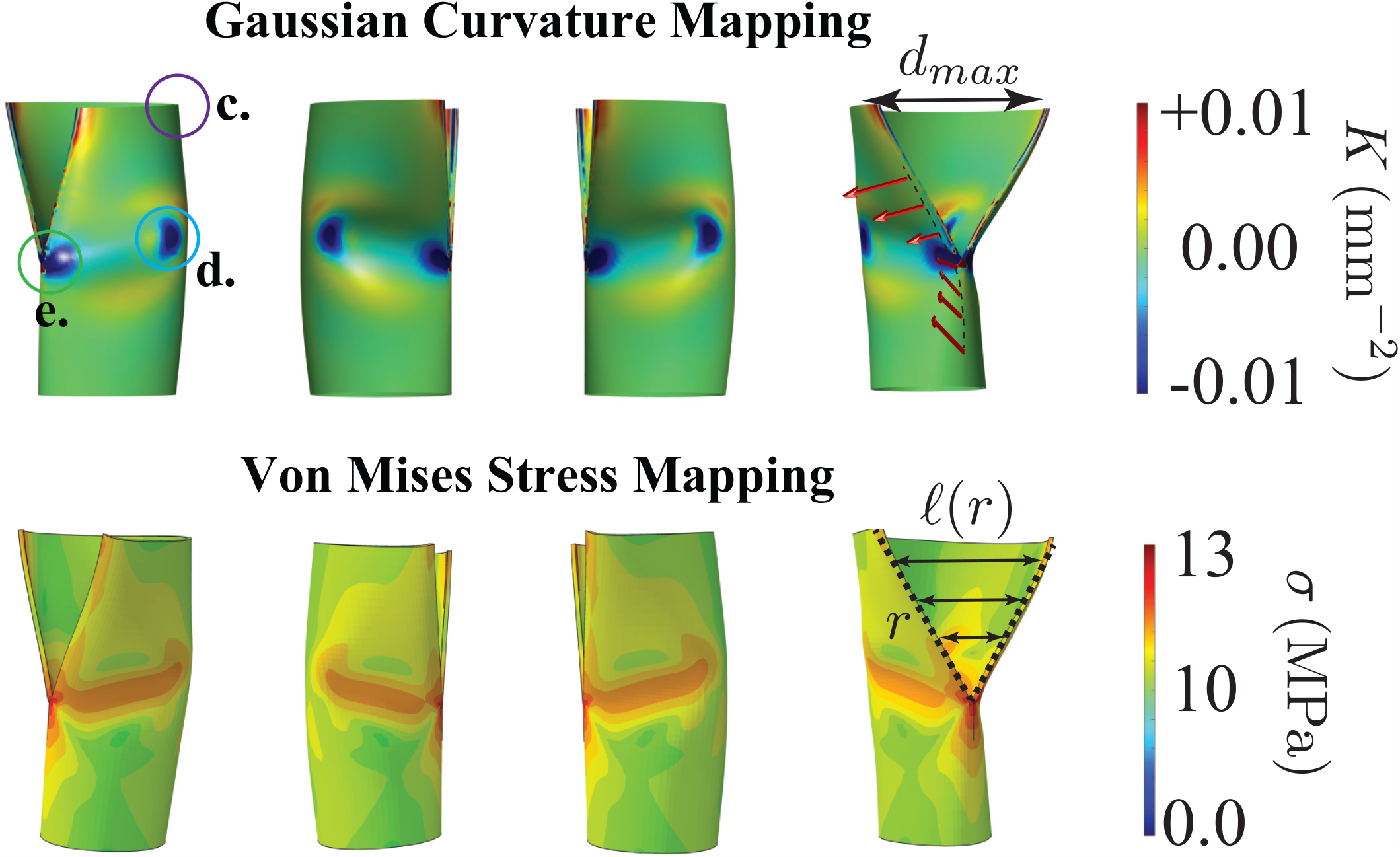
Representative maximally deformed (open to *d*_*max*_) geometries of the side-to-side anastomosis as simulated with FEM. The suture line is indicated on the bottom right figure in black dashed lines. The excess area created by pressurization has a triangular appearance and linear separation between the suture lines *£*(*r*) scales with radial distance *r* form the anastomotic vertex. The two principal curvatures, *κ*_1_ and *κ*_2_, are calculated at each point on the surface; Gaussian curvature, *K* = *κ*_1_ *κ*_2_, is mapped onto the deformed surface showing that the majority of the deformed geometry is isometric to the undeformed cylinder with *K* = 0 (green color). Four regions of curvature condensation are identified. Three of these correspond to the conical singularities discussed in figure 2: green circle around the anastomotic vertex (e.) is an *e*-cone, blue circle around remote region far-field from suture line (d.) is a *d*-cone, and a regular cone will appear at the end in vicinity of the purple circle (c.). There is curvature condensation in a fourth region along the suture line away from the vertex, especially towards the free edge, this is not further explored in this paper. The red arrows indicate the surface normals along a director in the undeformed cylinder, it shows how the *e*-cone functions as a pivot point for the normals. The *e*-cone charge is proportional to the separation between the suture lines, *£*(*r*) (see figure 2). Bottom row shows the same deformed geometry with the von Mises stress field mapped onto the surface. Of note, the stress is very low in all regions which deformed isometrically. However, the stress is two orders of magnitude above the baseline stress in regions corresponding to non-zero *K*, especially at the *e*-cone, *d*-cone, and ridge connecting them. This indicates that the observed stress focusing is strongly linked to the underlying geometry; supporting the hypothesis that the geometric singularities are driving the stress focusing.

Figure 3 outlines the basic geometric structure of a maximally open anastomosis using the deformed shape from our simulation. As the pressure is increased, the suture lines separate increasing the geometric charge of the associated *e*-cone and with it the maximal distance between the suture lines *€*(*r*). The formation of an *e*-cone to accommodate increase in lumen cross section is not unique to this geometry. HM strictureplasty (see Appendix Supplemental Figure 10), size-mismatched end-to-end (Billroth I) and the end-to-side anastomosis all include an *e*-cone along their main suture line (Appendix Figure 9 - with FEM of computed tomography derived rubber models). However, in the Billroth-I as well as in the side-to-side anastomosis, the unpressurized anastomosed tissue may assume a flat, folded relaxed state that does not show the induced *d*-cones. The normals to the tissue on each of the sides of the flat folded configuration are all approximately parallel. In circular cylinders, the normal are parallel only along straight lines that are oriented along the cylinder’s axis. Along the circumferential direction, the normals vary and complete a full turn around the cylinder. In particular, at any given point on a cylinder the normals vary only along one direction. In contrast, for a regular cone or an *e*-cone all normals meet at a single vertex (at which the normal is ill-defined). If we now form an *e*-cone on the side of a cylinder, there are regions in the vicinity of the *e*-cone where the normals vary also along the long axis of the cylinder. Normals in smooth developable surfaces can never cross and only meet at singular vertices. Upon pressurization of the anastomosis the effective *e*-cone opens and the limbs assume their cylindrical form far away from the suture. Consequently, there is a mismatch in local tissue orientation, which is reconciled by the formation of the *d*-cones.

At the vicinity of the vertex of a conical structure in infinitely thin (isometric) sheets, the normal curvatures may become arbitrarily large. Physical tissue is naturally associated with a finite thickness which regularizes this attempted divergence by compromising the underlying geometry and introducing in-plane strains. The diverging curvature along sharp creases that connect two conical structures are also similarly regularized. Geometrically, the in-plane deformations may give rise to Gaussian curvature, which typically vanishes for cylindrical surfaces. Mechanically, we expect these regions to experience significant in-plane stress, that could eventually affect tissue remodeling. Figure 3 depicts the emergence of Gaussian curvature and the stress-focusing in the vicinity of the conical structures and along the ridges connecting them. Bottom panel of figure 3 shows the stress distribution of the deformed shapes in the top panel; the transitional geometry shows a substantial increase in stress compared to the isometric cylindrical and conical regions. Comparing the top and bottom rows one can visually appreciate that non-zero *K*_*g*_ (geometric singularities) are linked to stress focusing.

### Tissue Response to a *d*-cone

#### Submucosal Collagen Fiber Remodeling Model

The next focus of this study is to explore the mechanism driving changes in the dispersion of submucosal collagen fibers in the above experiments. To elucidate this mechanism, we start the simulations with an aligned fiber state where the dispersion *κ* is set to zero. The remodeling of the submucosal collagen fibers is triggered in the regions where the stress becomes larger than a critical value: *σ > σ*_*cr*_, where we use the von Mises stress for an effective measure of the stress state (19). Within regions of collagen fiber remodeling, the orientations of the collagen fibers tend to gradually align with the major principal loading directions (20). For uniaxial loading this implies an alignment with the loading direction and hence a reduction in collagen fiber dispersion *κ*→ 0. In contrast, for equal biaxial loading where all directions are equivalent this leads to a more isotropically dispersed state where *κ*→1*/*3 (20). While such fiber reorientation models were applied to cardiovascular tissues, we present here the first integration of collagen fiber remodeling with the OGH constitutive law and enrich it to incoporate the complex stress-focusing and loading states induced by side-to-side anastomosis constructions. Specifically, the two major principal loading directions (hoop and longitudinal) dictate the preferable dispersion 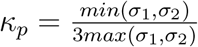 of the collagen fibers where 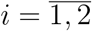 are associated with the element’ stress (stretch) state. The remodeling happens to adjust the initial fiber dispersion of each element towards the targeted preferable value. A remodeling rate equation (20) is used to govern this adjustment process: 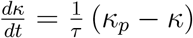 where *τ* is the rate of the remodeling which can be adjusted to capture how fast or slow the remodeling happens in real experiments.

The above model is implemented using the user material subroutine VUMAT in Abaqus (see Methods for details). Our simulations show that as the anastomosis opens and *d*-cones occur, remodeling happens in the stress-focusing regions (Figure 4F). The loading states in the remodeling zones are close to an equal biaxial stress state that drives the submucosal collagen fibers to experience a more dispersed state: *κ*_*p*_→1*/*3. Figure 4E shows this increase of the *κ* value from the initial dispersion state. The distribution of the dispersion values from the numerical simulations at the end of the remodeling process is qualitatively consistent with the experimental measurement shown in Figure 4B at POD 14. It is also noted in the simulations that as a result of this readjusmtent of the collagen fibers, the stress of the elements are alleviated in the remodeling area.

**Fig. 4.**
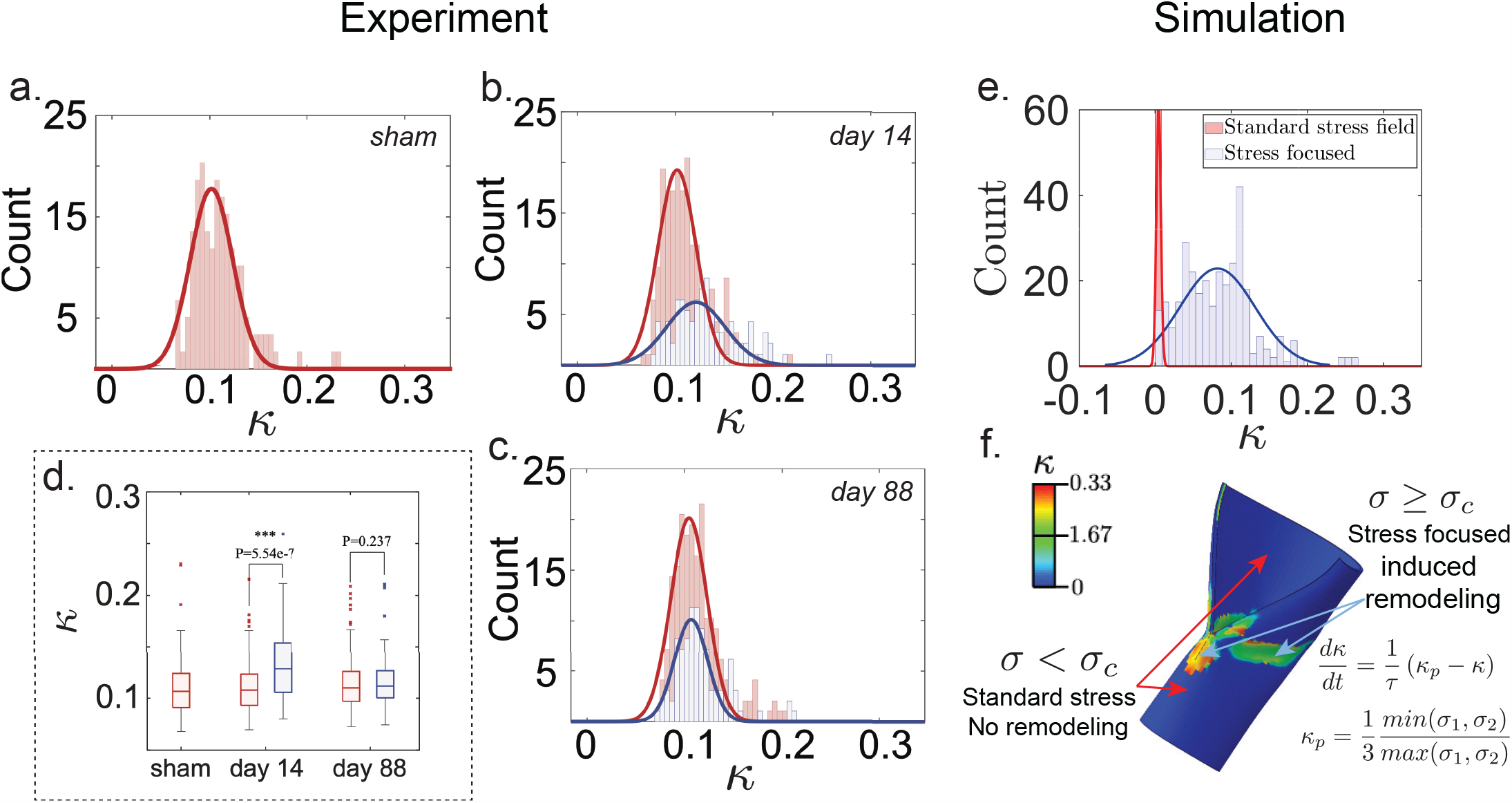
Distributions of *κ* values in (a.) sham operated mice and in (b.) and (c.) mice status post side-to-side anastomoses. In (a.), the distribution is from a normal segment of small bowel and has a single symmetric peak centered at *κ* = 0.11 indicating weakly dispersed oriented submucosal collagen fibers. In (b.) and (c.), the distributions are from the peri-anastomotic segments of bowel. Two populations of post-operative mice are analyzed: (b.) mice sacrificed on POD 14 and (c.) those sacrificed on POD 88. The vertex *e*-cone serves as the common origin of every side-to-side anastomosis. Following the method outlined in figure 5, the distributions in (b.) and (c.) are separated into two populations: near-*e*-cone (blue) and far-field (red). In (b.), the distribution becomes asymmetric with a shoulder appearing at the higher *κ* that can be fit to a separate distribution centered at *κ* = 0.13 containing only near-*e*-cone submucosa segments. The far-field region at POD 14 and both regions at POD 88 remain centered at *κ* = 0.11. (d.) shows the statistical difference between the near-*e*-cone and far-field regions which exists at day POD 14 is gone at POD 88; moreover, in all cases the far-field regions are indistinguishable form normal (sham operated) bowel. (e.) shows the output of a simulated side-to-side anastomosis using an OGH collagen reinforced material with stress-based remodeling. (f.) shows a representative output of post-remodeling *κ* values projected as a heat map onto the deformed geometry (one open limb of the anastomosis). The remodeling equations are solved only in finite elements that meet the stress criterium *σ* ≥ *σ*_*cr*_, with *σ*_*cr*_ a tunable control parameter. The simulations show that the stress criterium is only satisfied in the near-*e*-cone region of the anastomosis as seen in (f.), which is in agreement with the co-localization of geometric singularities and stress focusing seen in figure 3. For simplicity, the un-deformed collagen fibers are perfectly oriented so initially *κ* = 0. The remodeling rate equation evolves the per-element *κ* value to a target *κ*_*p*_ that is given as a normalized ratio of the finite element’s principle stretches. Consequently, isotropic biaxial principal stress/stretch leads to *κ* approaching 1*/*3 while anisotropic stress/stretch leads to *κ* approaching 0. (e.) shows that in non-stress focused regions of the anastomosis, where *σ < σ*_*c*_, *κ* remains 0; while in regions with stress focusing the distribution of *κ* values broadens, similar to the distribution broadening seen experimentally at POD 14 (b.).

#### Biological Experiments

The suture line is known to be an area of tissue injury and remodeling. However our interest is the biological response to the induced *d*-cones and Witten ridges away from the suture line; as such, a side-to-side anastomosis was performed in ten week-old male C57BL/6 mice who, following recovery were sacrificed on either post-operative day (POD) 14 or 88. Sham-operated mice (anesthesia, skin incision, and immediate closure) served as controls. In the experimental mice, anastomotic and peri-anastomotic tissues were harvested and sectioned. Images were then taken along the perimeter of the anastomosis away from the suture line every 0.2 mm, where tissue integrity was preserved such that Trichome staining allowed identification, visualization, and directional mapping of submucosal collagen fibers as a function of distance from the anastomotic vertex. Quantification of collagen fiber orientations was performed by fitting the fiber angles from each image to a transversely isotropic and *π*-periodic von Mises distribution. The mean collagen angle was approximately 30^*o*^ as reported in the literature (15). The spread of collagen angles per image is quantified through the dispersion parameter *κ* (see Methods). *κ* varies between 0 and 1/3 and is used in many continuum fiber reinforced models to take into account variance in fiber angles. Tissues with perfectly aligned fibers have *κ* = 0, while tissue with random fiber orientation have *κ* = 1*/*3. Figure 4A shows the distribution of *κ* values obtained every 0.2 mm along a segment of normal bowel in the control laparotimized animals with a mean of 0.11. Figure 4B shows similar data obtained in experimental animals at POD 14. The aggregate distribution in this case becomes asymmetric with a shoulder appearing at higher *κ* values that was not present in the control group. This shoulder can be understood by separating the different 0.2 mm segments into two groups based upon our FEM simulations and the predicted location of *d*-cones. The first group composes of images from tissue away from the anastomotic vertex and *e*-cone (red symmetric distribution in Figure 4B), while the second group is tissue forming the perimeter around the vertex (blue symmetric distribution in Figure 4B). The latter distribution (near-vertex) has a mean *κ* = 0.13 and is shifted to the right, while the former (far-field) has a mean of 0.11 similar to the control. Figure 4D shows that the far-field distribution is statistically identical to the controls, while the near vertex distribution is significantly different. When a similar analysis was performed on mice far from surgery at POD 88, no significant difference could be appreciated in the near-vertex and far-field groups, both being identical to the controls (Figure 4C).

While intestinal tissue is best modeled in its native state as a fiber-reinforced composite with two primary fiber-families running in a helical pattern through the submucosa in the *θ*−*z* plane (15), we observe a shift in the distribution of the orientation of submucosal collagen fibers in the immediate post-operative period. As noted in our POD 14 population, there is a distinct and statistically significant difference in the orientation of the submucosal collagen fibers when comparing those regions of bowel that experience standard stress fields with those that experience stress-focusing and *d*-cone deformation patterns, as informed by our FEM simulations in Figure 3. This can likely be attributed to the biological tissue’s desire to remodel in such a way so as to minimize the effects of these newly felt stress-focused fields. This shift in collagen fiber orientation leads the tissue away from its previous anisotropic response to stress to a more isotropic material, altering its deformation response to insults and likely its permanent function and appearance.

However, as post-operative time progresses, we note a return to the native orientation of these collagen fibers in those regions of the anastomosis that previously experienced stress-focusing and *d*-cone deformation patterns. This can likely be attributed to the permanent morphologic changes we observe in these anastomoses as they become grossly dilated and often fecalized at long time points. This adaptation of the affected bowel geometry via dilation leads to a relaxation in the stress-focusing initially felt after imposition of the non-linear geometry on the system, leading to re-organization of the submucosal collagen fibers into a conformation closely resembling its initial state. The scope of this work extends far beyond the bounds of this specific anastomotic construction.

We know that similar geometric aspects that lead to the formation of stress-focused points exist in other commonly used conformations as well. The strictureplasty, an operation performed to relieve narrowing of the lumen of the bowel, results similarly in the formation of *e*-cones through its efforts to longitudinally open and transversely close an incision through the anti-mesenteric surface of the bowel (9).

## Conclusions

While surgical reconstructions involve perturbations of both native geometries and soft tissue responses, these changes are often studied independently (9, 11, 13). This paper presents the first framework to elucidate the correlation between emergent geometric-focusing regions driven by the loaded reconstructed side-to-side anastomoses with the collagen fiber remodeling in bowel tissue. First, a novel operative technique to create a physiologically relevant small bowel side-to-side anastomosis is developed. This biological mouse model allows the characterization of submucosal collagen fibers and the spatial changes in dispersion parameter *κ*. Second, an FEM model of the opening of a side-to-side anastomosis is constructed which shows the emergence of conical singularities: *e*-cone at the suture line vertex and remote *d*-cones are induced far field in the undisturbed tissue.

Geometric analysis is utilized to explain the appearance and development of these geometric charges in the perturbed cylindrical bowel. The connection between FEM model and the experimental model allows the analysis of experimental data where two groups of data are revealed: the first one contains dispersion data from tissue around the vicinity of the *d*-cones and the second one contains dispersion data from tissue far away from these stress-focusing areas. The *κ* data shows a statistically significant increase in dispersion of collagen fibers in the first group as compared to the second one in the POD 14 while no statistically significant difference is observed in the sham-operated mice and for the long term POD 88. Since collagen fibers have been reported to be able to remodel according to the major principal loading directions (20), we hypothesized that how stress is distributed along the amorphous non-linear configuration of a side-to-side anastomosis is associated with the observed changes in the collagen fibers at POD 14. We confirmed this hypothesis by integrating a new stretch induced remodeling model coupled with OGH constitutive modeling incorporating criteria from previous experimental and numerical studies on collagen remodeling (20) into our FEM. The integrated experimental and computational framework provides better understanding of the interaction between geometry, mechanics, and biology occuring in surgicial anastomoses which is critical in the development of more scientific-based surgical manipulations to prevent unwanted tissue responses.

## Materials and Methods

### *In Vivo* Model

All mouse experiments were approved by the Institutional Animal Care and Use Committee (IACUC) of the University of Chicago (ACUP protocol 72417). Ten week-old male C57BL/6 mice (Charles River Laboratories©, Kingston, NY) were used for all experiments and were allowed unrestricted access to standard chow diet and tap water both pre- and post-operatively. All mouse experiments were run in duplicate.

All non-control or sham-operated mice were anesthetized and underwent laparotomy with handsewn small bowel anti-peristaltic side-to-side, functional end-to-end anastomosis using 8-0 monofilament, non-absorbable suture (Surgipro™ II Monofilament Polypropylene), following appropriate sterile technique. Laparotomy incision was closed in two layers using 3-0 braided, absorbable suture (Covidien Polysorb™). Anesthesia was obtained via intra-peritoneal injection of 100 mg/kg ketamine and 10 mg/kg xylazine (21). Post-operatively, all mice were resuscitated with 1 mL sterile water injected subcutaneously. Intra-operatively, a segment of small bowel was identified and transected, with special attention paid to ensure there was no interruption of blood supply to the bowel. The two ends of the bowel were spatulated along the anti-mesenteric surface and the common channel of the anti-peristaltic side-to-side, functional end-to-end anastomosis was created. The common anastomotic channel length was approximately 2-3 times the diameter of the bowel lumen 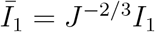, see Figure 5.

**Fig. 5.**
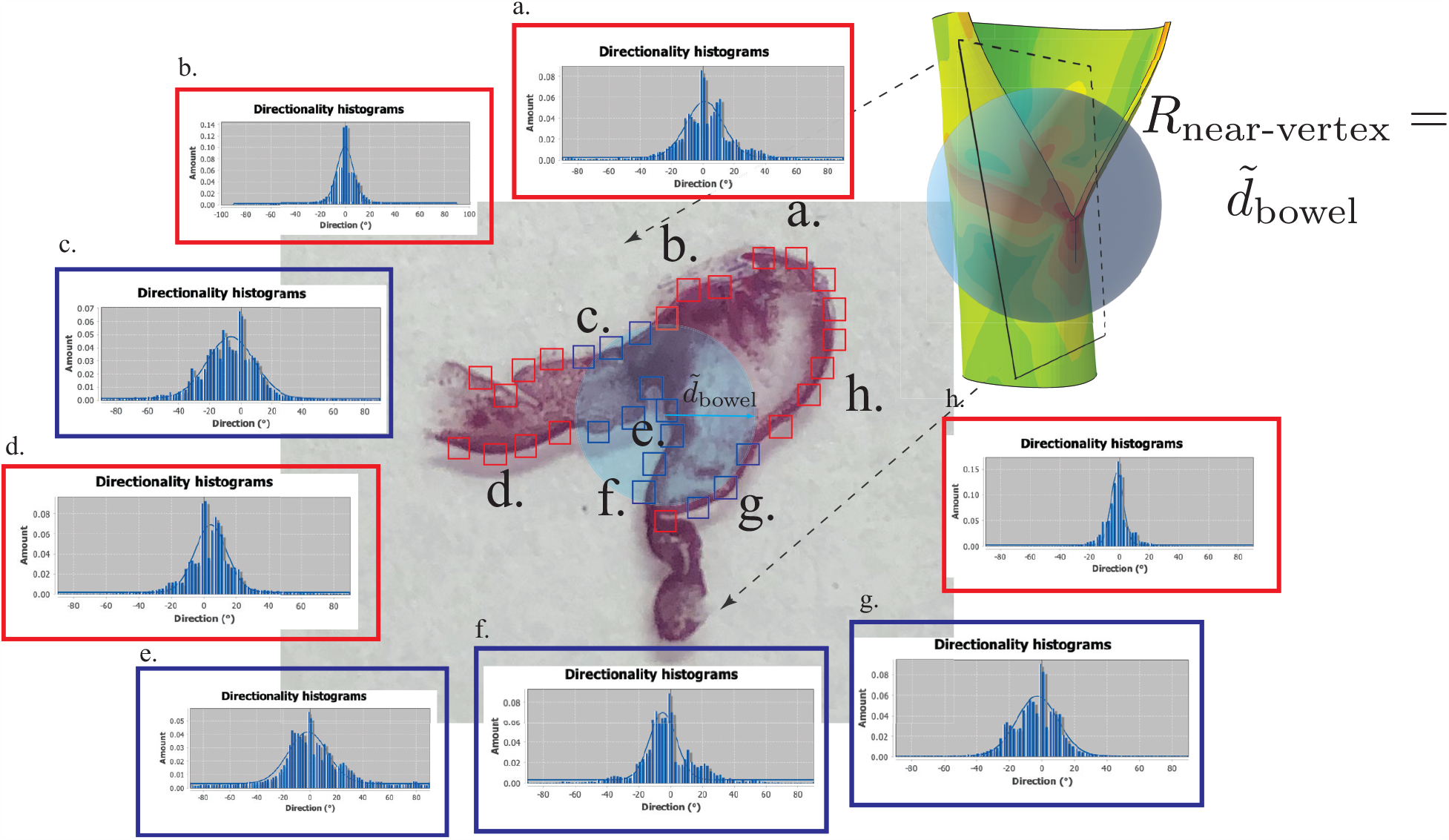
Representative microscopy section of side-to-side anastomosis (center). Both afferent (inflow) and efferent (outflow) limbs as well as the common channel were captured in sections used for subsequent analysis. High magnification region of interest (ROI images) were captured around the perimeter of each sample at predetermined locations, approximately every 0.2 mm of tissue (red and blue squares) taken around each anastomosis section, leading to ∼ 40 − 50 images (ROIs) per section. The angular distribution of the submucosal collagen fibers in each image (ROI) is measured. Representative distributions and their fit to equation 1 are shown a.-h. Also shown is the approximate mapping of a simulated bowel limb open as in an anastomosis and the microscopy image. ROI’s were partitioned based on their distance from the anastomotic vertex (*e*-cone). Two groups were generated: near-vertex group defined by ROIs located within a sphere centered at the vertex with radius 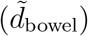 (blue squares) and a far-field group those outside of this spherical zone (red squares).

### Tissue Harvesting and Histology

The mice were sacrificed on either POD 14 or 88. The previous laparotomy incision was re-opened and the underlying bowel was carefully dissected out. The anastomosis and peri-anastomotic tissues were harvested and pressurized with 100 *μ*L of formalin to ensure bowel was fixed in a non-collapsed conformation. All harvested tissue was then formalin fixed and embedded in paraffin to prepare for sectioning and staining. Tissue samples were sliced into 5 *μ*m thick sections, which subsequently underwent Trichome staining. Sections were obtained at pre-determined depths, correlating with entry into the lumen of the bowel and areas of predicted stress-focusing as calculated using our FEM simulations. Figure 5 (center, low power 5x image) shows a representative section slice; both afferent (inflow) and efferent (outflow) limbs as well as the common channel were captured in sections used for subsequent analysis.

### Sample Imaging

All sample imaging was performed at 100x/1.3NA oil 0.2 mm WD magnification using a Zeiss Axioskop™ upright histology microscope. High magnification region of interest (ROI images) were captured around the perimeter of each sample at predetermined locations, approximately every 0.2 mm of tissue. Figure 5 shows the typical distribution of ROIs (red and blue squares) taken around each anastomosis section (∼ 40 − 50 ROIs per section).

### Submucosal Collagen Fiber Orientational Mapping

Initial image analysis was performed through Fiji and its Directionality plugin (v2.0). All captured microscopy images were cropped such that only the submucosal collagen fibers were included and rotated such that the horizon was set to *θ* = 0°. The aforementioned program was then implemented to characterize the angular orientation of submucosal collagen fibers in each ROI image via a Fourier transformation. These results were confirmed using a separately validated image processing algorithm (22). At each ROI image, the output was a probability distribution of the submucosal collagen angles centered at *θ* = 0° because of the original alignment of each image with the horizontal axis. Figure 5 (a-h) shows eight representative angular distribution plots (blue histograms) at different locations along the perimeter.

The normal collagen fibers of the small intestinal submucosa are known to consist of an interweaving lattice structure of two distinct fiber populations that run along the longitudinal axis of the bowel and intersect in a cross-ply fashion (15, 23). Further, it has been shown that the collagen fibers are distributed according to a transversely isotropic and *π*-periodic von Mises distribution, which is commonly used for directional data. We normalized the standard *π*-periodic von Mises distribution resulting in Equation 1, which is used to fit the angular distributions obtained at each ROI (see figure 5 ROI histograms and the overlaying solid blue curves which are the best fits to the histograms) (24).

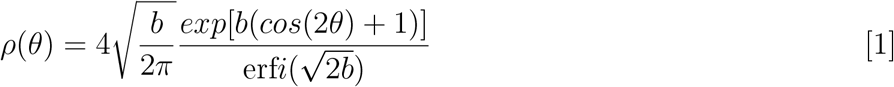

A widely used FEM of fiber-reinforced tissues is the OGH model (15, 25, 26). The non-uniform angular distribution of collagen fibers is captured in this model using a dispersion parameter *κ* which can be linked to the the concentration parameter *b* in Equation 1, using Equation 2 (16). At the end of our image analysis, each ROI of a given segment of bowel becomes associated with a single scalar value *κ*.

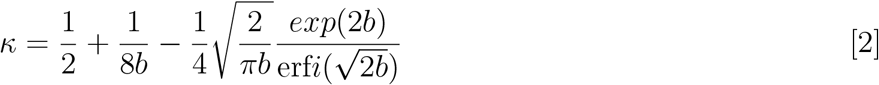

### Spatial Gradients and Distributions of *κ* Values

To study the effect of peri-anastomotic geometric singularities, ROI’s were partitioned based on their distance from the anastomotic vertex (*e*-cone). As shown in Figure 5, two groups were generated: ROIs located within a sphere centered at the vertex with radius ∼*d*∼_bowel_ (blue squares in Figure 5) and those outside of this spherical zone (red squares in Figure 5). The *κ* values for every image were then compared between these two groups using a one-tailed, two-sample unpaired t-test for samples of unequal variance (see Figure 4 for the distribution *κ* values within each group).

### Finite Element Modeling

A computational model for a side-to-side anastomosis was constructed in the commercial finite element (FE) software Abaqus (Dassault Systemes, MA) (27). The bowel is modeled as a cylinder with the realistic geometrical dimensions for small bowels used in the experiments (section A). Specifically, the diameter *d* ∼ 23*mm*, the thickness *t* ∼ 1*mm*, and the total length *L* ∼ 300 mm. A slit of length *l* = *L/*4 was created on the wall of a cylinder to model the suture lines on the anti-mesenteric surface of the tubular bowel. In the FE simulations, we model only one limb of the anastomosis, assuming symmetry with the second. This is done to decrease the computational cost of the simulation, since simulations with both limbs showed no appreciable differences compared to the single limb simulation (data not shown). Boundary conditions along the suture line are used to reproduce the mechanical effect of the kinematic coupling that the sutures impose. Pressure due to the flow of the luminal contents inside the bowel was applied to the internal surface to drive the opening of the anastomosis, *P*∼9 Pa. The main load bearing layer of the intestinal wall, the submucosa, is a fiber-reinforced layer with collagen distributed in the soft elastin matrix (15, 25, 26). To capture this structural aspect and the large deformation of the bowel tissue, the OGH model was utilized (15, 25, 26). We simulate the bowel wall as a single layer OGH model with the material properties adopted from previous work on intestinal mechanics (15): *μ* = 1.58 kPa, *k*_1_ = 22.01 kPa, *k*_2_ = 12.1, and *κ* = 0. The mean collagen fiber direction in the submucosa layer is *θ* = 30^0^ as suggested in (15). The model is based on a strain-energy function *W* = *W*_*m*_ + *W*_*f*_ composed of contributions from the elastin matrix *W*_*m*_ and the two families of collagen fibers *W*_*fi*_, where *i* = 1, 2. The specific forms for these energy functions are as follows:

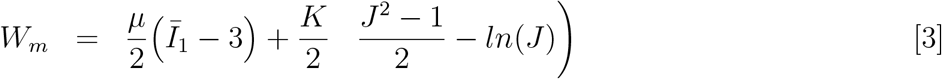

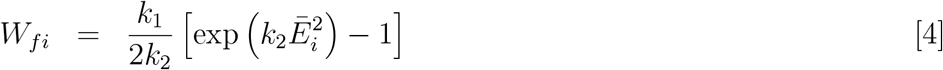

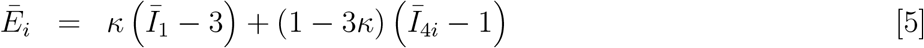

where μ and *K* are the shear and bulk moduli of the elastin matrix, respectively, *J* = *det*(*F*) represents the volume ratio between the deformed and undeformed tissue with *F* being the deformation gradient. To enforce a nearly incompressible behavior for the bowel material (15) where *J* → 1, a high numerical value for the ratio *K/μ* was used. 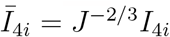 where *I*_1_ = *tr*(*C*) is the first invariant of the right Cauchy Green tensor *C* = *F*^*T*^ *F* widely used to study the nonlinear elasticity of soft elastomeric materials and tissues (15, 25, 26). Two symmetric families of collagen fibers *i* = 1, 2 are assumed to orient along an angle ±*θ* with respect to the circumferential direction, corresponding to two mean directions *A*^*i*^ (15). *k*_1_ is the fiber stiffness, and *k*_2_ is a dimensionless parameter correlating to the non-linearity of the collagen fiber’s response. *κ* is the dispersion of the collagen fibers. *κ* = 0 corresponds to an aligned state of collagen fibers along the mean direction and *κ* = 1*/*3 corresponds to an isotropic dispersion state. 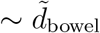 and *I*_4*i*_ = *A*^*i*^.*CA*^*i*^ gives the square of the stretch along the mean direction *A*^*i*^ of the *i*^*th*^ fiber family.

After the anastomosis opens and the loading is maintained, the stress state in *e*-cone and *d*-cone regions of the pressurized anastomosis induced the remodeling of the collagen fibers leading to changes in the fiber dispersion *κ*. To model this process, the preferable fiber dispersion will be updated as a function of the principal loading states in the bowel wall. While principal stresses can be used to trigger remodeling, we used here the principal stretches as these values potentially can be measured in experiments. Specifically, *κ*_*p*_ = *f* (*λ*_*i*_) where *λ*_*i*_, *i* = 1, 2 are the stretches along the major principal circumferential and longitudinal loading directions with *f* = *min*(*λ*_1_−1, *λ*_2_−1)*/*(3*max*(*λ*_1_−1, *λ*_2_−1)). Note that if |*λ*_1_| ≫ |*λ*_2_ |, the element experiences a near uniaxial loading state. Hence, the preferable value of *κ*_*p*_ becomes 0 as in the limit of uniaxial loading. In other words, the collagen fibers remodel toward the principal uniaxial loading direction, and thus exhibits a more aligned state. The same limit is obtained if |*λ*_2_| ≫ |*λ*_1_| . On the other hand, if |*λ*_1_| =|*λ*_2_|, the remodeling process in the limit of an equal biaxial loading in which *κ*_*p*_ becomes 1*/*3 is obtained. This means that collagen fibers becomes more dispersed and experience a more isotropic distribution. For any state of loading in between the preferable value of the dispersion can be determined. Collagen fibers tend to achieve this preferable dispersion value and the rate modeling equation as discussed in the main text is used to capture this process. This material model with the remodeling process is implemented through a material subroutine VUMAT. At each time increment, the principal stretch state for each element is obtained and used to compute *κ*_*p*_. For the elements that lie in the regions where the von Mises stress is greater than a critical value *σ > σ*_*cr*_, the rate remodeling equation is solved to update the fiber dispersion value. The obtained value for *κ*(*λ*_*i*_) is then used to update the stress in the element:

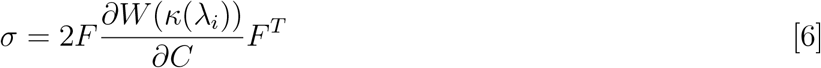

## Appendices

### *d*-cone Scaling: Comparison of Simple *d*-cone versus Remote Singularity in Model Side-to-Side Anastomosis

The remote singularity observed in the model side-to-side anastomosis has not been studied before. In Figure 2b, we make a geometric argument that this point must be a fully developable-*d*-cone in order to satisfy the global Gauss-Bonnet theorem. In Figure 3, we show that in the model side-to-side anastomosis this region carries a strong Gaussian curvature charge, where a strongly positive core is surrounded by a rim of strongly negative charge, such that the integrated Gaussian curvature is Φ = 0. Furthermore, the region containing the *d*-cone (spherical region with radius 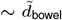 centered at the anastomotic vertex defined in Figure 5) has a complex non-homogeneous and increased stress distribution. The relationship between the spatial location of Gaussian curvature charge and the resulting stress field distribution can be highly non-linear, since the two are linked by a Laplace-type equation. The stress fields will also be sensitive to the details of the constitutive relationships used to model the material.

To more systematically study the stress distribution in this geometrically non-linear region, we performed simulations using a simpler neo-Hookean material, where the fiber stiffness *k*_1_ in the OGH model becomes zero, allowing comparison with existing theories of conical singularities. Figure 6 shows the evolution of the stress pattern as a function of decreasing shell thickness; for the thickest shell, a singular center of high stress appears far-field from the *e*-cone and is connected by a ridge like region of higher than baseline stress. As the thickness decreases, the far-field structure bifurcates and develops the Yoshimura buckling pattern typical for longitudinally loaded cylindrical shells consisting of crescent shaped singular points connected by elongated stressed regions. These are reminiscent of the well studied *d*-cone structures obtained by circular-confined thin flat sheets, and Lobokovsky-Witten stretching ridges that connect conical points in thin flat sheets (7, 29). Since there lacks a general theory of *d*-cones, we perform a comparison between the crescent stress distribution in the side-to-side remote singularity, with a standard method of generating a simple *d*-cone in plane sheets, indentation of a sheet into a cylinder (see Figure 7).

**Fig. 6.**
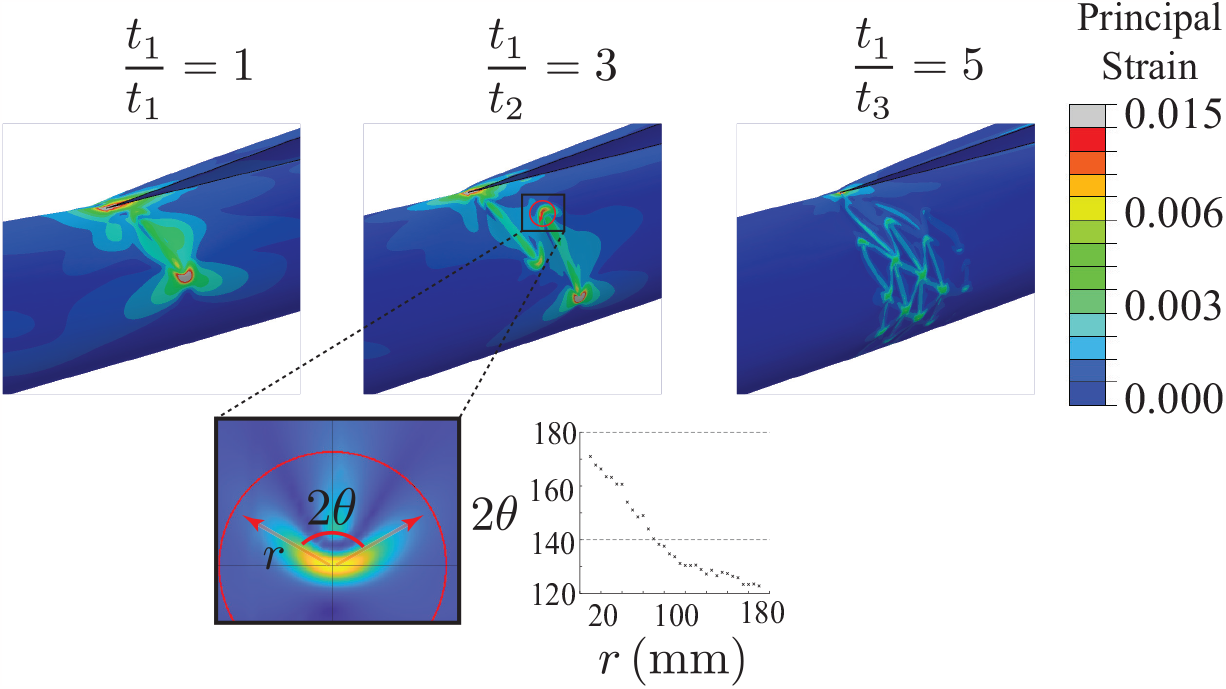
Three side-to-side deformed geometries with anastomosis open generating a vertex *e*-cone in cylinders of varying thickness: *t*_1_*/t*_2_ = 3 and *t*_1_*/t*_3_ = 5 are shown. To quantitatively study the far-field mechanical and geometric consequences of the *e*-cone, we employ a simplified homogeneous elastic material model in these simulations. In all three cases, all of the strain is again concentrated in the region of the transitional geometry as shown in Figure 3, where the biologically relevant non-linear fiber-reinforced OGH material model is used. In all three cases, the *e*-cone vertex carries a large degree of tensile strain in a similar distribution. Interestingly, the far-field high-strain structures evolve in a cascading self-similar pattern with the thickest cylinder having a single crescent-like focus of high strain which bifurcates into two daughter centers which again bifurcate leading to a pattern qualitatively very reminiscent of crumpling. In crumpling, centers of high strain driven by non-isometric deformation called *d*-cones are connected by weaker strain-focused lines termed stretching ridges or the Witten ridge (7, 29).

**Fig. 7.**
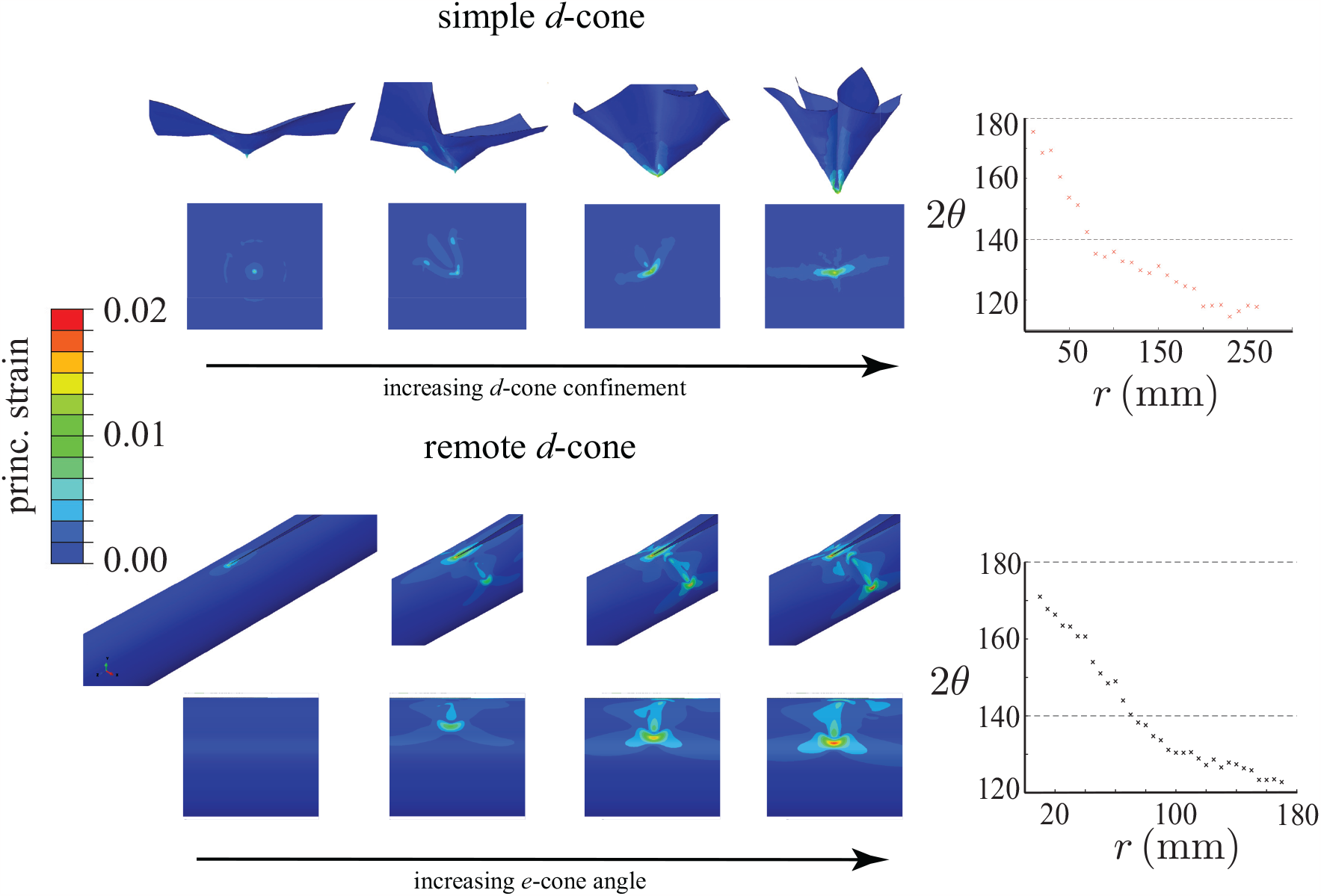
The evolution of the far-field strain pattern with thickness (Figure 6) qualitatively supports the hypothesis that this region of the cylinder deforms by forming *d*-cones locally. Given the lack of a general theory of *d*-cones, to prove that the remote singularity seen in our simulations is indeed a *d*-cone, we opted to compare the strain distribution of a simulated simple *d*-cone with the remote far-field *d*-cone. The simple *d*-cone is generated by indenting a flat membrane into a hollow cylindrical support, this is the classic loading geometry well known to generate a *d*-cone at the point of indentation (7, 18). We show both the deformed geometry and the projected strain distribution onto the undeformed initial shape. For the simple *d*-cone, the initial core region shows a symmetric strain distribution as expected for localized stretching due to indentation. Once the membrane undergoes a macroscopic bending/folding, the classic crescent-like distribution of strain at the *d*-cone core appears (18). With further confinement, the crescent distribution remains focused but evolves in its shape for an elastic system such as simulated here. The bottom row shows the evolution of a single remote *d*-cone on the cylinder with increasing *e*-cone angle; the same crescent shape is observed in the strain distribution around the core. This data shows that the simple *d*-cone and the remote *d*-cone have qualitatively the same distribution of strain in the core region, allowing us to make the claim that the far-field high strain singularity in the cylinder is indeed a *d*-cone. The opening angle, 2*θ*, of the crescent-like strain distribution seen in *d*-cones is measured as a function of radial distance away from the peak core stress as outlined in Figure 8. Note that the angle reaches an asymptotic limit once a given distance from the core itself. The simple *d*-cone shows a limiting 2*θ* ∼ 115^*o*^, while the remote *d*-cone has 2*θ* ∼ 125^*o*^. This proves quantitatively that the two structures are nearly identical and as such that the remote far-field strain focused singularity induced by the *e*-cone is a *d*-cone. Moreover, the opening angle falls into the theoretical range calculated for *d*-cones (18).

Top panel in Figure 7 shows the appearance of a simple *d*-cone as the elastic sheet evolves from a local symmetric stretching to global folding and local stress-focusing as the *d*-cone evolves. Bottom panel in Figure 7 shows the appearance and evolution of the remote high stress singularity in the side-to-side simulation. There is strong qualitative similarity between the appearance of the crescent-like strain distribution of the simple *d*-cone and the remote singularity in the cylinder. To allow for a quantitative comparison, we must measure and compare the opening angles of the crescent stress distributions (see inset of Figure 6). Our analysis is done at a fixed time step, far along into the simulation that we can safely assume a steady-state stress distribution has been attained. We import the (von Mises) stress data around the stress-focused region from Abaqus to Matlab (Mathworks, MA) using the package Abaqus2Matlab (30). This data is defined on the undeformed cylinder. For ease of viewing and subsequent analysis, we unroll and flatten the cylinder to a rectangle, using the standard mapping (*θ, z*) → (*x, y*), where *θ* is the azimuthal coordinate and *z* is the axial coordinate on the cylinder (see Figure 7). Finally, using peak detection, we isolate the point singularity we wish to study, and keeping in mind the measurements to follow, also interpolate this isolated data over a fine grid. We wish to measure quantities along the arc of the crescent singularity, i.e., as a function of the arc length coordinate *s* (see Figure 8). In other words, we wish to extract a meaningful 1-dimensional data field, say *σ*(*s*) for stress, from a 2-dimensional one, *σ*(*x, y*). To do this, we draw circles of increasing radius *r* centered at the tip of the crescent (located using peak detection algorithms), and measure *σ* only along this circular contour. Here, the radius *r* ≈ *s*, the desired arc length coordinate, since the crescent is not sharply curved. This gives a polar distribution of the data along the circle (as a function of the angle *φ*), which has a tri-lobular structure, as expected (each lobe is a maximum or peak), see Figure 8B. Since we are only interested

**Fig. 8.**
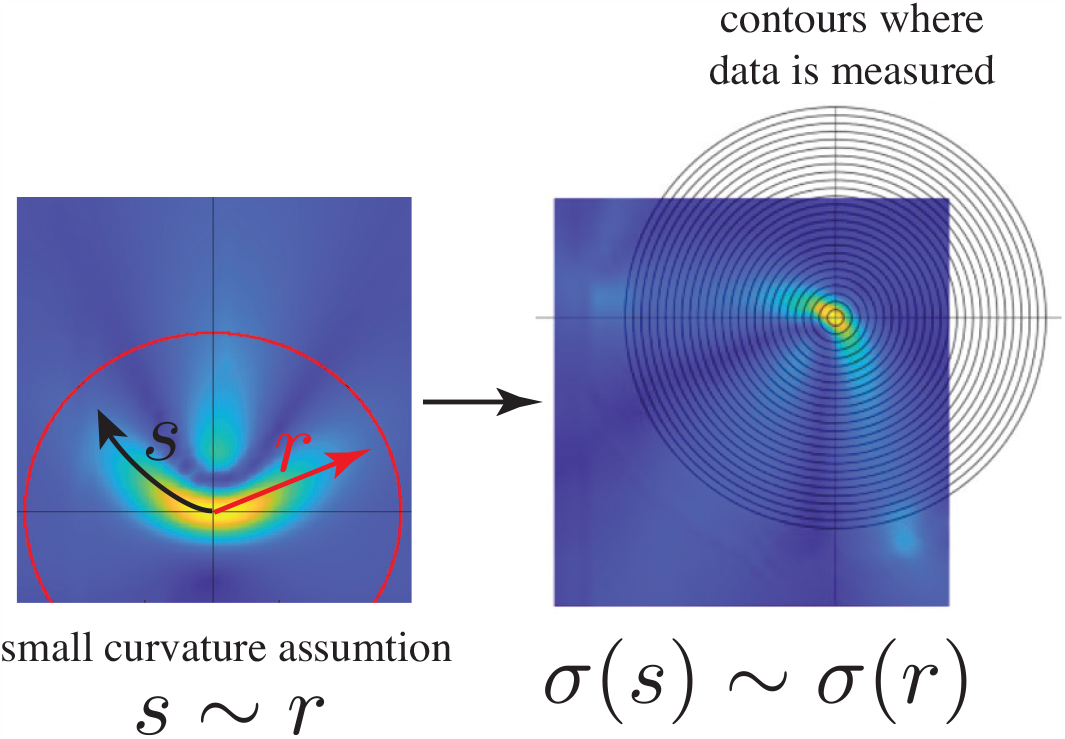
von Mises stress field projected onto a rectangular grid for extraction of angular information of stress dependence.

**Fig. 9.**
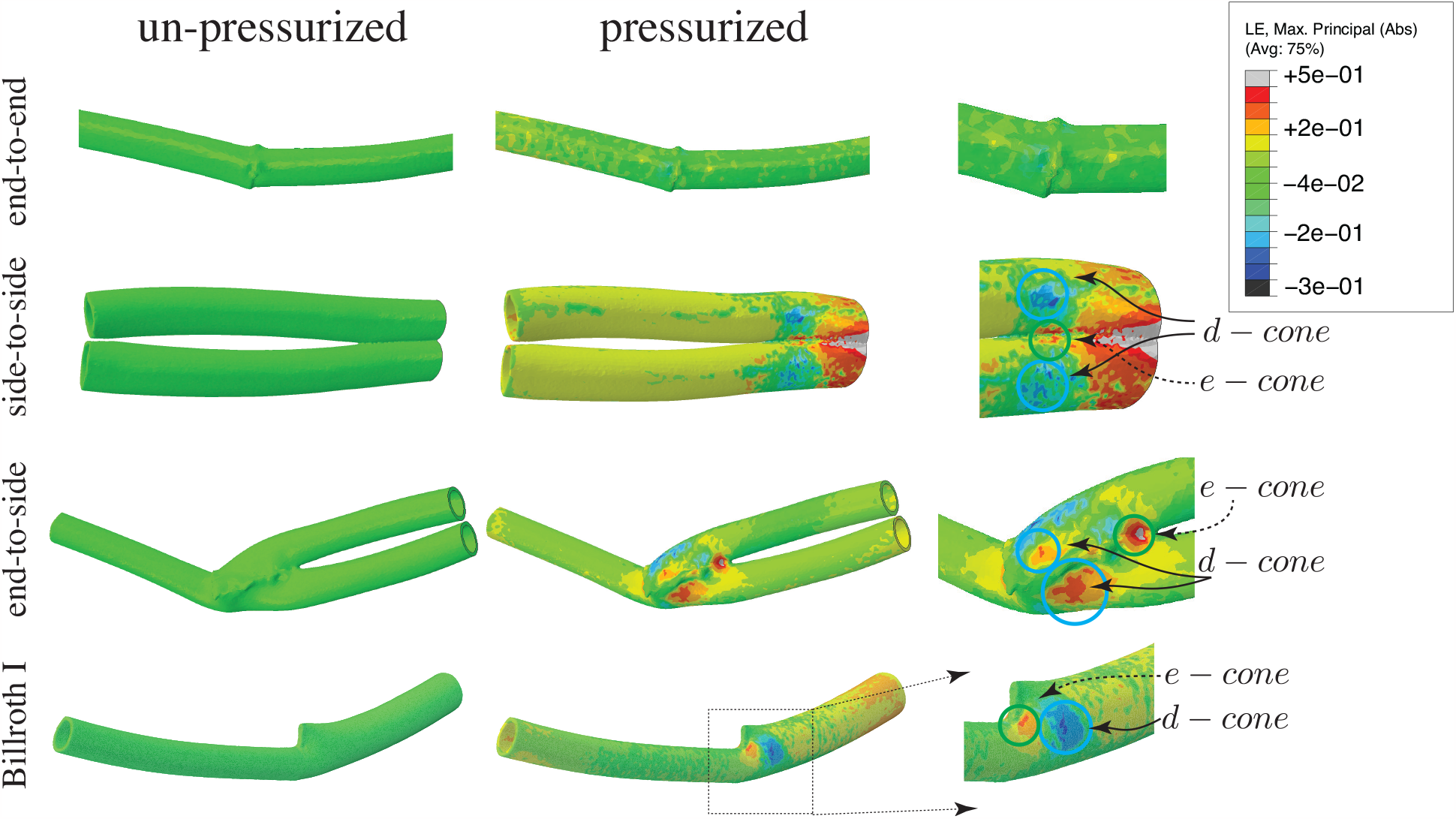
FEM simulations of CT-derived geometries of end-to-end, Billroth I, end-to-side, and side-to-side where *e*-cone and *d*-cones are shown to occur in the latter three anastomoses.

**Fig. 10.**
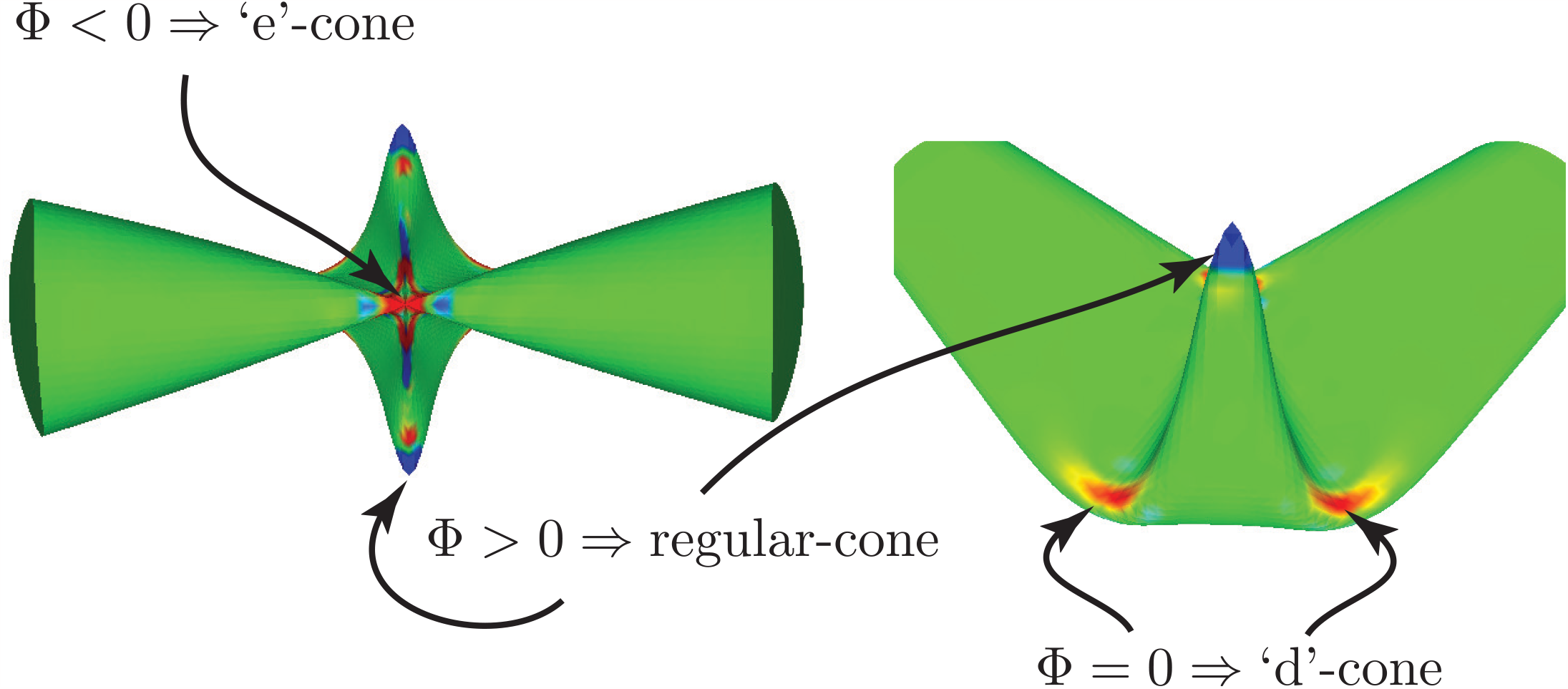
The geometry of conical points in a Heineke-Mikulicz strictureplasty. Our prior work (9, 11) showed that the shape is dominated by the ‘*e*’-cone which forms by joining the two vertices of the initial enterotomy during the transverse closure. The regular-cones flank the suture line and as noted in the main text are typically though of as dog ears. By the Gauss-Bonnet theorem, since a strictureplasty only involves making a cut and re-orienting the cut via transverse closure, the overall topology of the deformed cylinder is preserved: Φ_’e’-cone_ + Φ_regular-cone_ = 0. Away from the suture line four flanking ‘*d*’-cones appear.

in the crescent here, we only care about the two side peaks. Again, using peak detection, we find the angular locations of the three lobes. Shifting the central peak to the y-axis (*φ*= 0), we can then measure the opening angle of the two side lobes, call them *φ*_1_ and *φ*_2_. Measurements show that *φ*_1_ = −*φ*_2_ = *φ*_*cresc*_. This gives us a value for that particular circle, of radius *r*. Doing this for a range of circles, we get the desired distribution *φ*_*cresc*_(*r*) ≈ *φ*_*cresc*_(*s*) along the arc of the crescent. The *d*-cone opening angle is Δ*θ* = 2*φ*_*cresc*_. Similarly, we can get the stress along the two arms of the crescent, *σ*_1_(*s*) and *σ*_2_(*s*). Measurements also show that *σ*_1_(*s*) ≈ *σ*_2_(*s*). The same analysis applies to the simple *d*-cone situation, with just the initial data preparation being simpler since the undeformed geometry is already a rectangle, and not a cylinder.

Figure 7 shows our data for the canonical *d*-cone opening angle 2*θ* plotted as a function of distance from the peak stress. The simplest theory of *d*-cones states that 2*θ* is constant at 140^0^ (17), though this is only in the limit of linear elasticity and planar initial geometries. The nearly identical dependence of 2*θ* on radial distance for the two simulations strongly supports the hypothesis that the remote singularity being driven by the *e*-cone is indeed a *d*-cone.

### Model Surgical Anastomoses

Figure 1 schematically shows multiple anastomoses, including an end-to-end with cut bowel perimeters matched, an end-to-end with mismatched perimeter lengths (Billroth I), an end-to-side, and a side-to-side. In this section, we show that all of these anastomotic geometries, except the end-to-end, when pressurized show the appearance of remote singularities. Figure 9 shows FEM simulations beginning with an unpressurized stress-free geometry. *e*-cones and regular cones exist even in the stress-free state, however, what is most striking is the strong appearance of *d*-cones upon uniform pressurization. The side-to-side geometry was extensively studied in the main text. Here we show that *d*-cone avoidance can be very difficult, moreover *d*-cones can appear spontaneously even when the stress-free configuration already has a fixed *e*-cone such as the end-to-side and Billroth I.

## ACKNOWLEDGMENTS

Please include your acknowledgments here, set in a single paragraph. Please do not include any acknowledgments in the Supporting Information, or anywhere else in the manuscript.

## Notes

No conflicts of interest.

### Competing Interest Statement

The authors have declared no competing interest.

### Summary of Updates

Updated figures and addition of new simulation data for anastomotic geometries.

